# Failed Metabolic Adaptation to Stress Underlies Pathogenesis in a Heterozygous Genetic Disorder

**DOI:** 10.1101/2025.04.13.648535

**Authors:** Robert Harmon, Erin McCarthy, Annette Krol, Jonas Braun, Aashutosh Shetti, Lisa Godsel, Ziyou Ren, Bethany Perez-White, Samuel Weinberg, Cory L. Simpson, Russell Dahl, Yogesh Goyal, Murali Prakriya, Eran Cohen-Barak, Roni Dodiuk-Gad, Amy S. Paller, Kathleen Green

**Affiliations:** Department of Pathology; Department of Dermatology; Department of Pediatrics; Skin Biology and Diseases Resource-based Center; Department of Cell and Developmental Biology; Department of Medicine; Department of Pharmacology Feinberg School of Medicine, Northwestern University, Chicago, Illinois, USA; Robert H. Lurie Comprehensive Cancer Center, Northwestern University, Chicago, IL USA; Institute for Stem Cell and Regenerative Medicine, University of Washington, Seattle, Washington, USA; Neurodon Corporation, 9800 Connecticut Drive, Crown Point, IN, 46307, USA; Center for Synthetic Biology, Northwestern University, Chicago IL, USA; CZ Biohub Chicago, LLC, Chicago, IL, USA; Department of Dermatology, Emek Medical Center, Afula, Israel; Bruce and Ruth Rappaport Faculty of Medicine, Technion, Haifa, Israel; Department of Medicine, University of Toronto, Toronto, Ontario, Canada; Lewis-Sigler Institute for Integrative Genomics, Princeton University, Princeton, NJ 08544, USA

## Abstract

Disorders linked to heterozygous variants occupy a continuum in terms of the timing and severity of phenotypic emergence. An important question regarding this variability entails the effect stress has on the residual protein function. Using Darier disease (DD), caused by heterozygous variants of the SERCA2 calcium pump, as a model, we uncovered a potential connection between extrinsic stress and pathogenesis. The skin lesions characteristic of DD entail loss of intercellular adhesion and rarely appear pre-adolescence, suggesting that factors beyond heterozygosity contribute to disease pathogenesis. Testing whether age-related stressors contribute to DD, we show that DD patient-derived keratinocytes subjected to stress yield twice the reactive oxygen species of controls, accompanied by greater disruption of intercellular adhesion. Metabolic analysis of DD cells revealed perturbation of the pentose phosphate pathway (PPP), a stress response system responsible for regenerating antioxidants like glutathione. At baseline, DD cells had less free glutathione but an increase in protective glutathione-based modifications of SERCA2, a reversible form of protein oxidation. With stress, DD cells form an aberrant, heavily glutathionylated perinuclear halo consisting of keratin and the intercellular adhesion component, desmoplakin. We propose a model whereby SERCA2 heterozygosity causes mild oxidative stress that under homeostatic conditions can be buffered by glutathionylation. When stressed, the depleted glutathione store is shunted towards the desmoplakin-intermediate filament system at the expense of SERCA2, rendering it vulnerable to damage. A lesional flare, then, would represent a case of more complete SERCA2 inhibition and a novel example of how heterozygous disorders interact with stress to disrupt intercellular adhesion.

## Introduction

The interaction of heterozygosity with stress underlies many carcinogenic phenomena. Application of the concept to heterozygous genetic disorders, particularly those with unpredictable onset and severity, may also prove useful. To explore this possibility, we take advantage of a genetic skin disorder, Darier disease (DD), linked to heterozygous variants of the gene encoding the endoplasmic reticulum (ER) calcium pump SERCA2, *ATP2A2* [1]. Ion transfer across membranes represents a primordial cellular activity [2, 3]. Nonetheless, despite our fundamental understanding of widely expressed ion channels and pumps, application of this knowledge disproportionately focuses upon cardiac and neuronal function [4]. One such scenario entails the rhythmic movement of calcium from cardiomyocyte cytosol to the sarcoplasmic reticulum by SERCA2, an ATPase-driven pump required for maintenance of cardiac function [5-7]. Illustrative of the untapped potential of such knowledge, SERCA2 is widely expressed and implicated in numerous pathologies, including diabetes, yet the intermediates connecting pump function to an array of disease endpoints often remain unclear [8-11].

DD belongs to a group of skin disorders in which keratinocytes of the stratified, epidermal tissue fail to adhere to one another, a condition called acantholysis, leading to blistering and infection [12, 13]. DD and the related disorder, Hailey-Hailey disease (HHD), linked to pathogenic variants in *ATP2C1*, both lack targeted therapeutics [14]. Typically, DD arises from heterozygous mutations in *ATP2A2* resulting in either reduced SERCA2 expression or enzymatic activity, suggestive of haploinsufficiency as the primary pathogenic mechanism [15]. Historically two schools of thought have emerged to explain the disruption of intercellular adhesion in DD. One focuses on the perturbation of cytosolic calcium resulting in interference with the desmosome-intermediate filament network [16], and the other on diminished lumenal ER calcium stores, resulting in ER stress [17, 18]. While efforts to target the disorder therapeutically by normalizing ER calcium or indirectly through intermediary pathways linking mutant SERCA2 to skin deterioration have been published [19, 20], they have yet to reach widespread use in the clinic.

The present study takes a new approach towards understanding pathomechanisms associated with SERCA2 deficiency. Patients born with heterozygous SERCA2 mutations do not exhibit skin defects at birth. Most develop acantholytic lesions in the second or third decade of life and do so within specific anatomical sites (so-called seborrheic or oily areas), often in response to stressors including heat, UV irradiation, psychological stress and cutaneous super-infection [13]. These observations suggest that heterozygosity sensitizes intercellular adhesions to exogenous stressors.

Here we model extrinsic stress with the oxidative stressor, menadione, and show that DD keratinocyte adhesions exhibit heightened sensitivity to the compound. The effect abates upon treatment with the widely used antioxidant, N-acetylcysteine (NAC). Disruption of cell adhesion correlates with an inability to dampen ROS levels in response to menadione treatment, consistent with a muted antioxidant response. Metabolomic analysis of DD cells reveals perturbation of the pentose phosphate pathway (PPP), a metabolic pathway parallel to glycolysis that produces the NADPH necessary to regenerate the antioxidant activity of glutathione. We propose a model whereby SERCA2 heterozygosity causes PPP perturbation and mild oxidative stress initially buffered by glutathionylation, a process by which glutathione binds to and protects protein from irreversible oxidation. However, under exogenous oxidative stress, the already-depleted GSH store is shunted towards an abundant perinuclear pool of junctional proteins at the expense of SERCA2, rendering it vulnerable to oxidative damage. Our study provides a key conceptual advance in the study of genetic disorder onset: pathogenic phenotypes driven by heterozygous genotypes can stay latent and emerge through conditions mimicking “homozygous” loss-of-function conditions. In the case of DD, we hypothesize that SERCA2 haploinsufficiency endangers the residual wildtype copy by rendering it prone to oxidation and inactivation. The onset of DD cutaneous flares, generally not present at birth, would equate to intermittent unmasking of a phenotype by stress, essentially eliciting homozygous behavior from heterozygous cells.

## Results

### Modeling DD with cultured patient keratinocytes

Existing animal models of SERCA2 heterozygosity, at present, fail to mimic the pathognomonic epidermal acantholysis seen in DD [21]. To investigate the effect of SERCA2 haploinsufficiency on human keratinocytes, we use an in vitro model of cultured keratinocytes isolated from genetically characterized DD patients (DD1-4) or healthy adults (see Methods for donor information) (**Fig1A**). Western blotting reveals that patient cells express ∼70% of the SERCA2 produced by control cells, indicating that lost protein expression only partially accounts for subsequently observed defects (**Fig1B-C**). Calcium imaging via Fura-2 of the DD2 isolate indicates a roughly two-fold increase in baseline cytoplasmic calcium levels, aligning well with a 50% reduction in SERCA2-dependent ER calcium sequestration observed in prior studies (**Fig S1A-B**) [19]. Switching cells to media containing 2mM CaCl2 following inhibition of SERCA2 by thapsigargin typically elicits a large spike in Fura-2 signals arising from store operated calcium entry (SOCE) [22]. Of note, SOCE is hyperactivated in the DD2 cells compared to controls, perhaps implying that DD cells constitutively fail to support ER calcium sequestration and, thus, constantly activate SOCE channels (**Fig S1c)**. The expression and calcium imaging data supported further use of the cells as a model for SERCA2-dependent pathology.

**Figure 1.**
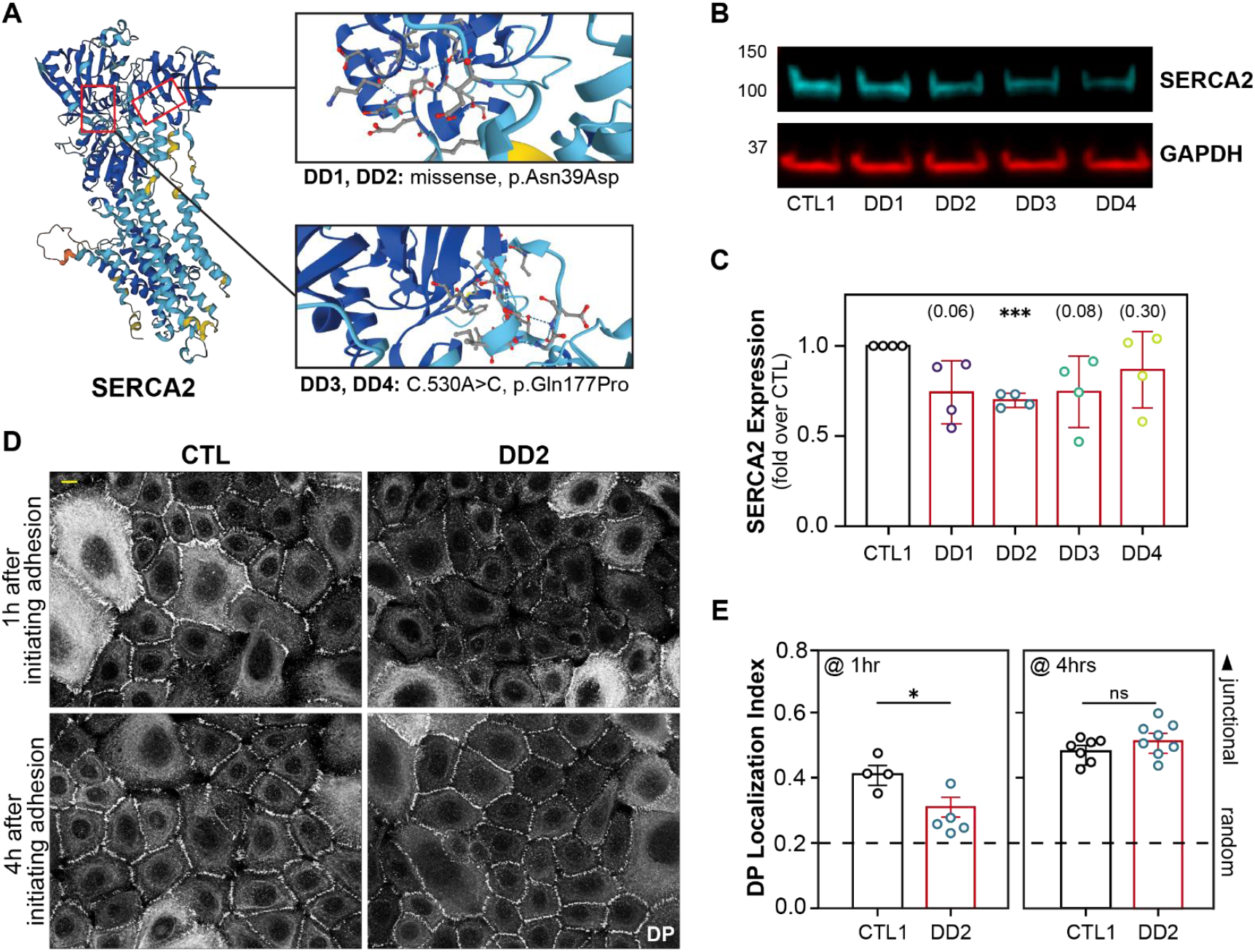
Keratinocytes isolated from DD patients model adhesion defects. a) Structural context of SERCA2 mutations present in tested samples. SERCA2 structure derived from AlphaFold prediction by EMBL-EBI [59, 60]. b) Western blot analysis of SERCA2 levels in DD cells compared to a control, adult keratinocyte isolate. c) Quantitation of western blot analysis. Values in parentheses represent p-values from one-sample t-tests (hypothetical mean=1, n=4 independent experiments). Bars equal mean relative quantity +/- s.e.m, circles represent values from each repeat. d) Immunostaining for DP, 1hr after switching cells from 0.07mM to 1.2mM CaCl2. scalebar= 10um e) Quantitation of DP border localization adhesion stain index (percent of cell boundaries occupied by pixels belonging to the top 20% with respect to DP intensity on cell by cell basis, see **Fig S2**). Values are presented for cells 1hr and 4hr post calcium switch. Bars equal mean relative quantity +/- s.e.m, circles represent values from each repeat.

To study cell-cell junction maturation, intercellular adhesion was induced by raising the media [Ca2+] from 0.07mM to 1.2mM [23]. Compared to controls, DD keratinocytes show significantly diminished accumulation of DP at nascent adhesions 1hr after initiating cell junction formation. This defect is readily detected by our custom pipeline incorporating Cellpose-based image analyses designed to assess the extent to which DP localizes to the adhesive interface between cells (**Fig S2 and Methods**) [24, 25]. As DD arises from heterozygosity, residual SERCA2 and downstream adhesion maturation should respond positively to addition of SERCA activators. Indeed, the activator, CDN1163, developed by Kang et al [9] completely reverses DP accumulation defects (**Fig S2**). This indicates that desmosome assembly is directly and reversibly impacted by loss of SERCA2 activity in our model, rather than other cellular damage inflicted by long-term calcium dysregulation. We also observe that, in culture, junction assembly normalizes in DD cells with time, suggesting impairment but not full blockade of desmosome assembly. Four hours after calcium switch, control and DD keratinocytes exhibit comparable DP localization (**Fig1D-E**). This suggests that acantholytic flares are likely precipitated by external factors that exceed the compensatory capacity of heterozygous keratinocytes.

### DD keratinocyte adhesion is hypersensitive to oxidative stress due to antioxidant dysfunction

Patients with DD experience cutaneous flares in response to environmental factors, most of which share the downstream effect of eliciting oxidative stress. We hypothesized that application of a stressor would exacerbate the observed defect in cell-cell junction assembly and better recapitulate a fully acantholytic phenotype. To elicit oxidative stress, we treat cells with menadione, which drives overproduction of mitochondrial reactive oxygen species (ROS) [26]. In our 2D model system, control cells exposed to menadione for 4h establish robust desmosomal plaques. DD cells, however, localize the majority of DP to a halo around the nucleus, far from its proper localization at cell-cell junctions. This defect is reversible via addition of the antioxidant n-acetylcysteine (NAC) (**Fig2A-B**). Formation of a perinuclear keratin ring has been observed in previous studies of DD and is reminiscent of keratin localization in keratinocytes from the blistering disease epidermolysis bullosa (harboring keratin mutations) when exposed to mechanical stress [27, 28].

**Figure 2.**
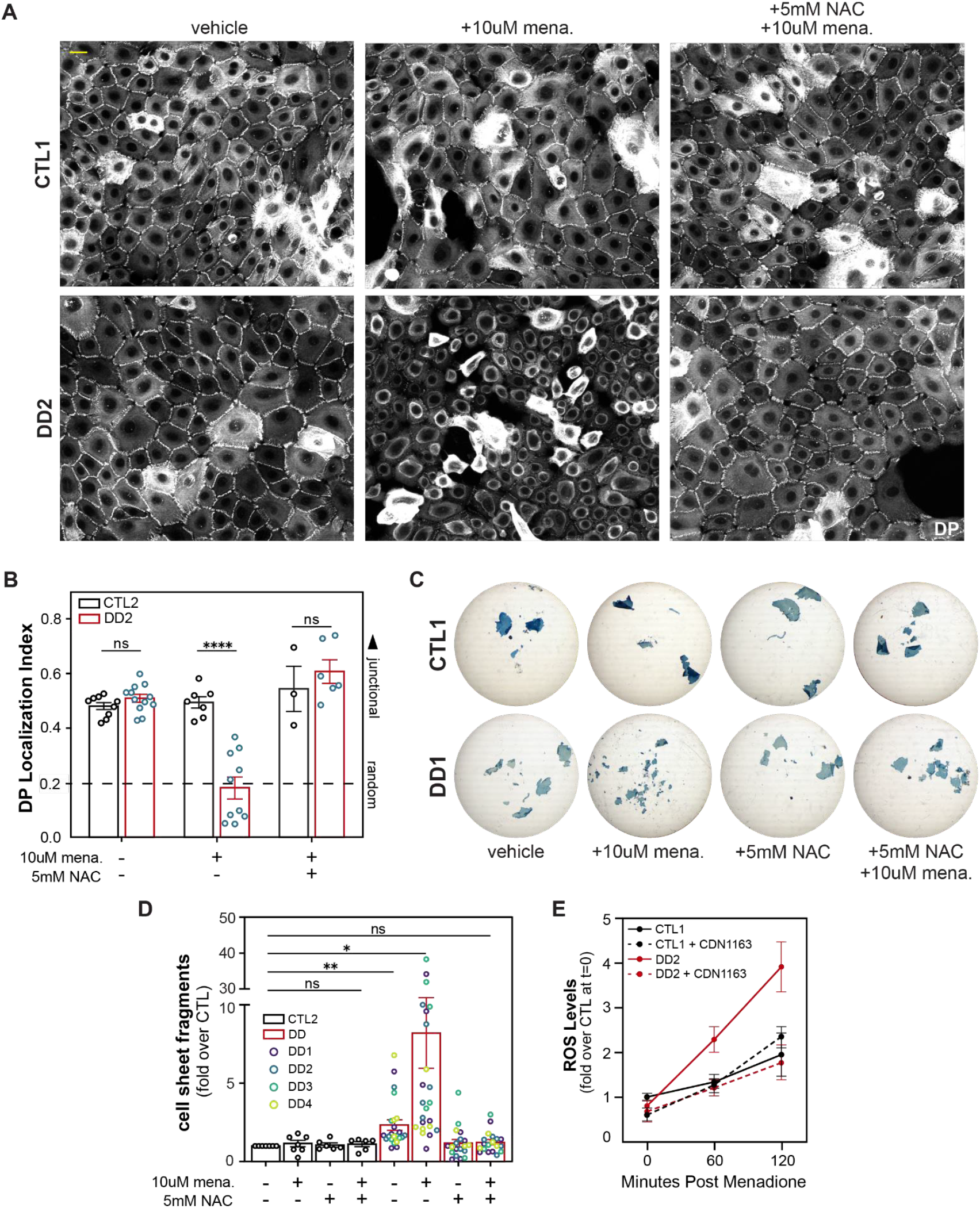
DD Keratinocyte adhesion is hypersensitive to oxidative stress arising from dysfunction in antioxidant response. a) Immunostaining for DP, 4hrs after switching cells from 0.07mM to 1.2mM CaCl2 with or without menadione (mena) or NAC. scalebar = 25um b) Quantitation of DP localization adhesion stain index; p-values derived from two-tailed t-test. Bars equal mean relative quantity +/- s.e.m of 8 independent experiments, circles represent values from each repeat. c) Dispase assay 4hrs after CaCl2 switch and addition of menadione, NAC, or both, showing decreased epithelial sheet integrity in DD cells treated with menadione (mena). d) Quantitation performed by counting the number of fragments in each well of a 6-well plate after physical agitation. n=4-7; p-values derived from two-tailed t-tests. e) ROS levels were measured using a CellRox assay; values displayed represent raw values normalized to CTL at t=0. Individual points represent an average of n=3 experiments for each condition +/- s.e.m.

Dispase assays enable functional assessment of intercellular adhesive strength and entail enzymatically removing an intact cell monolayer from the substrate, subjecting it to mechanical agitation, and inferring adhesion strength as an inverse relationship with the extent of cell sheet fragmentation [29]. We observe in four isolates from three patients that DD cell sheets have a mild, but statistically significant, adhesion defect at baseline. Consistent with staining results, the addition of menadione greatly exacerbates this phenotype. In both cases, NAC reverses the defect, supporting the therapeutic potential of antioxidants for DD (**Fig2C-D**). The ability of an antioxidant to correct the baseline adhesion defect suggests that DD cells possess an inherent deficiency of redox capacity that can directly impact adhesion strength, particularly under oxidative conditions.

To determine what aspect of redox control is altered in DD, we measure ROS levels using a CellRox assay and note no difference in ROS levels between the disease and control cells at baseline. Upon addition of menadione, DD cells do not dampen the resultant increase in ROS as efficiently as controls (**Fig2E**). Of note, the phenotype is reversible upon addition of the SERCA activator, CDN1163. This result serves as confirmation that SERCA2 deficiency disrupts the capacity of cells to detoxify ROS in keratinocytes. Taken with the correction of baseline adhesion by NAC, these data indicate a possible perturbation of the core metabolic pathways and/or antioxidant capacity of these cells.

### SERCA2 deficiency disrupts the Pentose Phosphate Pathway (PPP) and metabolic stress response

To investigate how SERCA2-dependent calcium dynamics impact the metabolic response of keratinocytes to stress, we conducted untargeted metabolomic analysis of control and DD cells with and without menadione. Comparison of controls to DD cells in the absence of menadione reveals 29 metabolites that differed significantly (p<0.05) and an additional 16 with notable trends that did not reach statistical significance (p<0.1) (**Fig3A**). Pathway enrichment analysis indicates that intermediates of the pentose phosphate pathway (PPP) and glutathione metabolism are among those most significantly altered in DD keratinocytes (**Fig3B**), driven by diminished levels of 6-phosphogluconic acid (6-PG), allocystathionine, glutathione, glutamate and a ribose species referred to hereafter as R5P (ribose-1-phosphate, ribose 5-phosphate, ribulose 5-phosphate are indistinguishable by the instrument). The PPP is a pathway, parallel to glycolysis, which facilitates cellular responses to stress by producing nucleotide precursors required for DNA repair and the NADPH needed to replenish the antioxidant activity of glutathione. Of relevance to the observed sensitivity of DD cells to oxidative stress, the active/reduced form of glutathione (GSH) is diminished in DD isolates. This baseline deficiency likely contributes to the sensitivity of DD cells to ROS induced by menadione.

In pairwise comparisons, each individual DD isolate clusters distinctly from CTL1 in partial least squares discriminant analyses, demonstrating a consistent deviation of DD metabolics from that of a normal keratinocyte culture (**Fig S3A**). R5P was the only specific metabolite to be significantly downregulated (p<0.05) in each of the four DD isolates individually, pointing to the likely importance of PPP insufficiency in DD pathogenesis (**Fig S3B**). This is further demonstrated by correlation analysis of control and DD responses to oxidative stress. When CTL1 keratinocytes are exposed to menadione, a set of 44 metabolites show strong positive Pearson correlations (rho= 0.74 +/- 0.15; mean+/- sd), matched by a negatively correlated group of 27 metabolites (rho= -0.48 +/- 0.23 sd). A coordinated loss of glycolytic-TCA cycle products and a surge of PPP metabolites drives these metrics, consistent with prior studies of stress responses in A549 epithelial cells (**Fig3C-E, Fig S3C**) [30]. These homeostatic metabolic adaptations collapse in DD cells (rho= 0.27+/-0.42 and -0.08 +/- 0.31, respectively). Baseline glutathione deficiencies of ∼40%, corroborated by a plate reader assay, are retained even after the precipitous decrease elicited by menadione (**Fig 3F, Fig S3D**). DD cells likely fail to shunt intermediates from glycolysis to the parallel PPP resulting in diminished capacity for regenerating GSH and an inadequate antioxidant defense (**Fig 3G**). We hypothesize that, at baseline, these metabolic abnormalities elicit measurable GSH deficits but do so at levels that only modestly affect cell adhesion, consistent with what would be predicted for the buffering of mild oxidative stress by oxidation of GSH and consumption of PPP intermediates. However, these baseline deficits become problematic when stressors place heightened antioxidant demand upon already weakened response systems, which likely precipitates the full acantholytic phenotype in active DD lesions.

**Figure 3.**
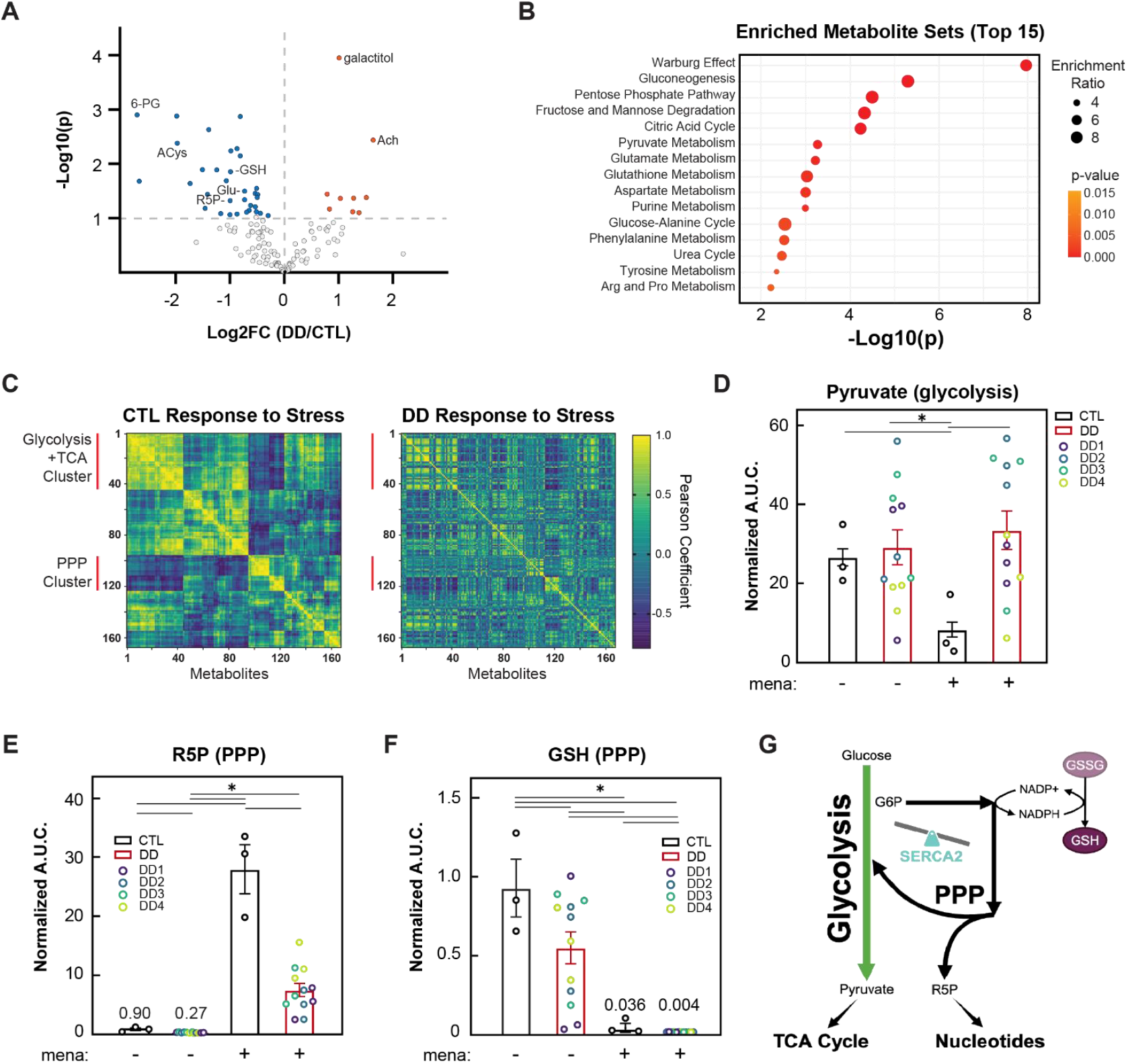
The metabolic response to stress is aberrant in DD keratinocytes. a) Volcano plot demonstrating baseline metabolite differences in DD versus CTL keratinocytes. Samples were collected for 3 independent experiments. Each metabolite was normalized to the median area under the curve (AUC) obtained from LC/MS analysis. Individual metabolites which differed in a two-tailed t-test with equal variance at p<0.1 are color coded. Blue and red dots represent metabolites that were diminished or elevated, respectively, in DD samples compared to controls. b) Enrichment analysis of significantly altered metabolites demonstrating overrepresented metabolic processes. c) Correlation analysis derived from pairwise comparisons of no stress versus menadione-treated values for the following isolates: CTL1 and DD1-4. Yellow indicates metabolites which either decreased or increased together in response to stress. Blue represents metabolite pairs which change in opposite directions. Panel on left contains clusters determined for the CTL1 sample. Map on right represents CTL1 clusters applied to the DD metabolite response to stress (correlations determined after calculating average metabolite changes across DD1-4). Samples were normalized to median AUC from 3 independent experiments prior to clustering. d-f) Response of pyruvate, R5P (D-ribose-1/5-phosphate / D-ribulose 5-phosphate), and GSH (glutathione-reduced form) to menadione. Samples normalized to median AUC. Columns represent mean control and DD AUCs after normalization +/- s.e.m. Asterixes indicate p<0.05 in paired t-tests of values paired within each independent experiment. g) Diagram of relationship between salient metabolic pathways.

### Stressed DD keratinocytes glutathionylate a perinuclear protein pool including DP

The canonical model of glutathione cycling indicates that, when oxidized, GSH dimerizes via disulfide bonding to an inactive form, GSSG. This transformation can be reversed via glutathione reductase and NADPH supplied by the PPP [31]. However, GSH can also be spent in the process of protein glutathionylation. Under oxidative conditions, GSH forms disulfide bonds with reactive protein cysteines. The process is thought to be a protective mechanism since, although indicative of oxidative stress, glutathionylation is reversible [32]. In the absence of glutathionylation, however, cysteines can become irreversibly oxidized, typically resulting in protein inactivation. SERCA2, itself, is known to be sensitive to oxidation and a known target of glutathionylation [33]. We hypothesized that our ROS and glutathione measurements may be consistent with low levels of oxidative stress that are buffered at baseline by transferring free glutathione to a glutathionylated pool of protein.

To address this idea, immunostaining for the glutathione adduct enables visualization of global glutathionylation, with and without stress. As seen previously, DP collapses around the nucleus in DD cells under stress. The cytoskeletal element anchored at desmosomes in basal keratinocytes, Keratin-14 (K14), also collapses with DP into a perinuclear ring. This aberrantly localized shield of K14 and DP colocalizes with the signal for glutathionylated protein (**Fig 4A**). This suggests that these critical elements may well act as a sink for free glutathione in stressed, disease cells, and DP glutathionylation may be a unique and crucial response of DD keratinocytes to oxidative stress.

**Figure 4.**
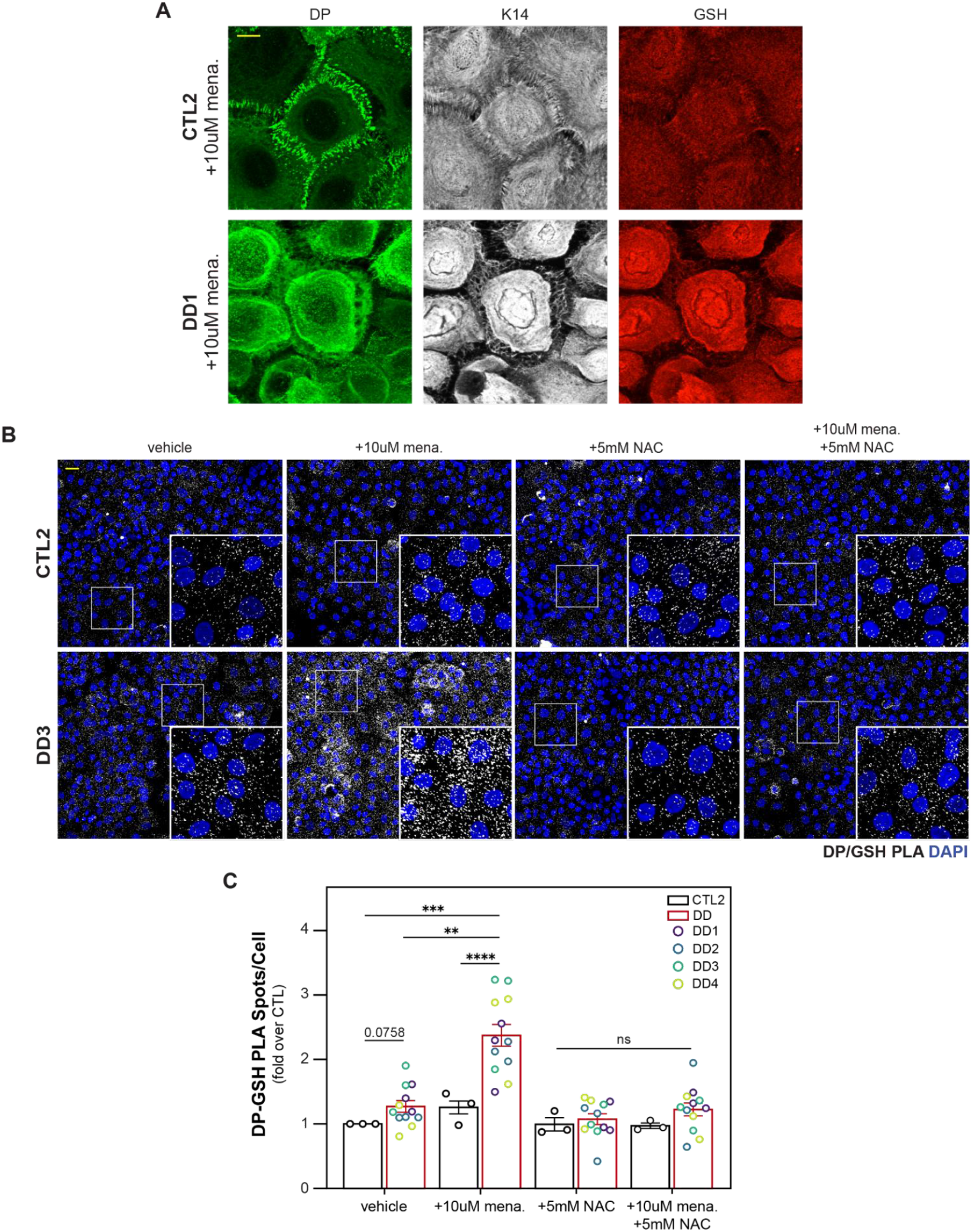
Stressed DD keratinocytes display an abundant perinuclear pool of glutathionylated protein including keratin and DP. a) Immunostaining for indicated markers in control and DD keratinocytes 4h after switching cultures to 1.2mM CaCl2 in the presence of 10uM menadione (mena). scalebar=10um b) Proximity ligation assay (PLA) imaging of sites where DP and glutathionylation antigens are in close proximity, 4 hrs after raising CaCl2 concentration in the presence of indicated treatments. scalebar = 25um. c) Quantitation of PLA signals normalized to vehicle treated control measurments, n=3 independent experiments, asterix indicate p<0.05 in t-test comparisons (one-sample test for comparison to vehicle treated controls, two-tailed equal variance t-tests for additional comparisons).

To more accurately assess the glutathionylation status of DP, we utilized a proximity ligation assay (PLA). In this assay, antibodies target two features of interest, in our case glutathionylated cysteines and DP. If the two targets are in close proximity (within ∼40nm), a PCR-based reaction produces a discrete fluorescent signal in situ. When applied to three sets of patient cells, this approach reveals no significant difference in DP glutathionylation between control and DD cells at baseline. However, under oxidative stress induced by menadione, DD keratinocytes have significantly higher levels of DP-proximal glutathionylation, an effect that is reversible with NAC (**Fig4B-C**). Together with our metabolomic findings, these data suggest a defect in the ability of DD cells to regenerate reduced GSH, with protein glutathionylation contributing to the mechanism of this phenotype.

### Stress-induced SERCA2 inactivation and loss of cell-cell adhesion via impaired glutathione buffering

The importance of glutathione in DD is supported by previous work showing that SERCA2, itself, becomes glutathionylated in response to oxidative stress [33, 34]. Though indicative of progression towards complete and irreversible oxidation, the modification initially protects and, potentially, elevates SERCA2 activity. We hypothesize that glutathionylation buffers the mild, baseline oxidative stress caused by SERCA2 heterozygosity, directly protecting the residual WT SERCA2 copy from irreversible oxidative damage and possibly increasing activity to compensate for haploinsufficiency.

To address this possibility, we assessed the glutathionylation status of SERCA2 via PLA and observed that DD keratinocytes have significantly higher levels of SERCA2-proximal glutathionylation than controls in the absence of exogenously applied stress (**Fig5A-B**). Both the antioxidant NAC and a SERCA2 activator reverse the effect, indicating that its own elevated glutathionylation/oxidation likely traces back to deficient SERCA2 activity associated with heterozygosity. Thus, in DD keratinocytes, SERCA2 is constitutively susceptible to oxidation at baseline, albeit in a reversible manner.

**Figure 5.**
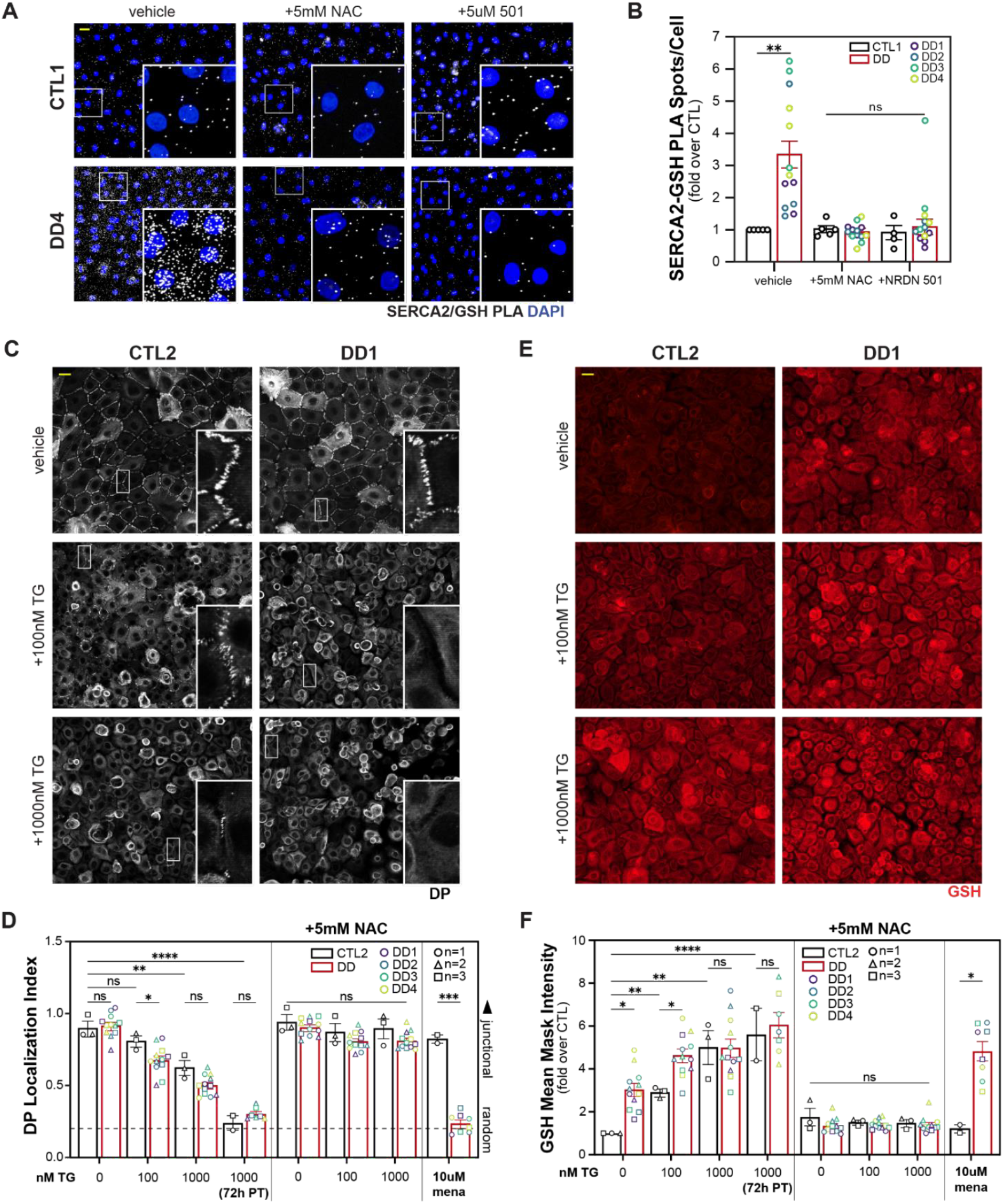
SERCA2 is partially oxidized in DD cells at baseline and total SERCA2 inhibition mimics menadione treatment. a) PLA imaging for sites of close proximity between SERCA2 and glutathionylation antigens 4hr after raising cells to 1.2mM in the presence of indicated treatments. b) Quantitation of SERCA/glutathionylation PLA with treatment of SERCA activator (NRDN-501, 10uM) and NAC. PLA signals normalized to vehicle treated controls, n=5 independent experiments, asterix indicate p<0.05 in test comparisons (one-sample tests for comparisons to vehicle treated controls, two-tailed equal variance t-tests for other comparisons). c) Immunostaining for DP 4hr after raising cells to 1.2mM CaCl2. Cells were pretreated with thapsigargin (TG) and indicated concentrations for 1hr prior to raising calcium, with the exception of the bottom panels which were pre-treated overnight. TG was kept in media during the calcium switch. d) DP adhesion localization indices for indicated treatments, n=3 independent experiments, asterix indicates p<0.05 for two-tailed t-test comparisons between controls and DD isolates. e) Immunostaining for glutathionylated proteins 4hrs after a calcium switch. Administration of TG and calcium was carried out as in panel c. f) Mean signal intensity for global protein glutathionylation normalized to vehicle treated control cells under the indicated treatments, n=3 experiments, asterix indicate p<0.05 (one-sample t-tests for comparisons to the vehicle treated control and two-tailed equal variance t-tests for other comparisons). scalebars=25um.

To complement observations using DD keratinocytes we assessed the impact on glutathionylation of thapsigargin (TG), a SERCA inhibitor. Immunostaining of desmosomal components in CTL cells treated with TG are remarkably similar to DD cells treated with menadione (**Fig5C-D**). Though previously unreported, TG also elicits a marked increase in global protein glutathionylation (**Fig5E-F**). Consistent with haploinsufficiency, both adhesion and glutathionylation responses in DD cells occur at lower TG concentrations than controls. Treatment with NAC blocks TG-dependent effects on both glutathionylation and adhesion, suggesting that glutathionylation links SERCA2 to intercellular adhesion (**Fig5C-F**). The data suggest that fully acantholytic DD lesions likely represent regions of near total SERCA2 inactivation, with exogenous stressors mimicking the effects of TG.

Considering our glutathionylation, metabolic, and adhesion data, we propose a model of stress sensitivity in DD in which SERCA2 heterozygosity compromises cellular metabolism to diminish PPP-dependent support of the antioxidant defense system (**Fig 6**). The modest oxidative stress that results is readily buffered by the large pool of GSH which remains. However, upon exposure to stress, the system fails to mount an adequate antioxidant response, the remaining GSH store is redistributed to glutathionylate a large perinuclear pool, including the keratin cytoskeleton, and less concentrated proteins such as SERCA2 are left unprotected, prone to irreversible oxidation and inactivation. The feed forward mechanism leads to disintegratation of cell adhesions, reminiscent of TG-treatment, and the acantholytic flares characteristic of DD. More broadly, this study highlights the importance of considering the metabolic aspect of those maladies, with their own characteristic endpoints, linked to SERCA2 deficiency.

**Figure 6.**
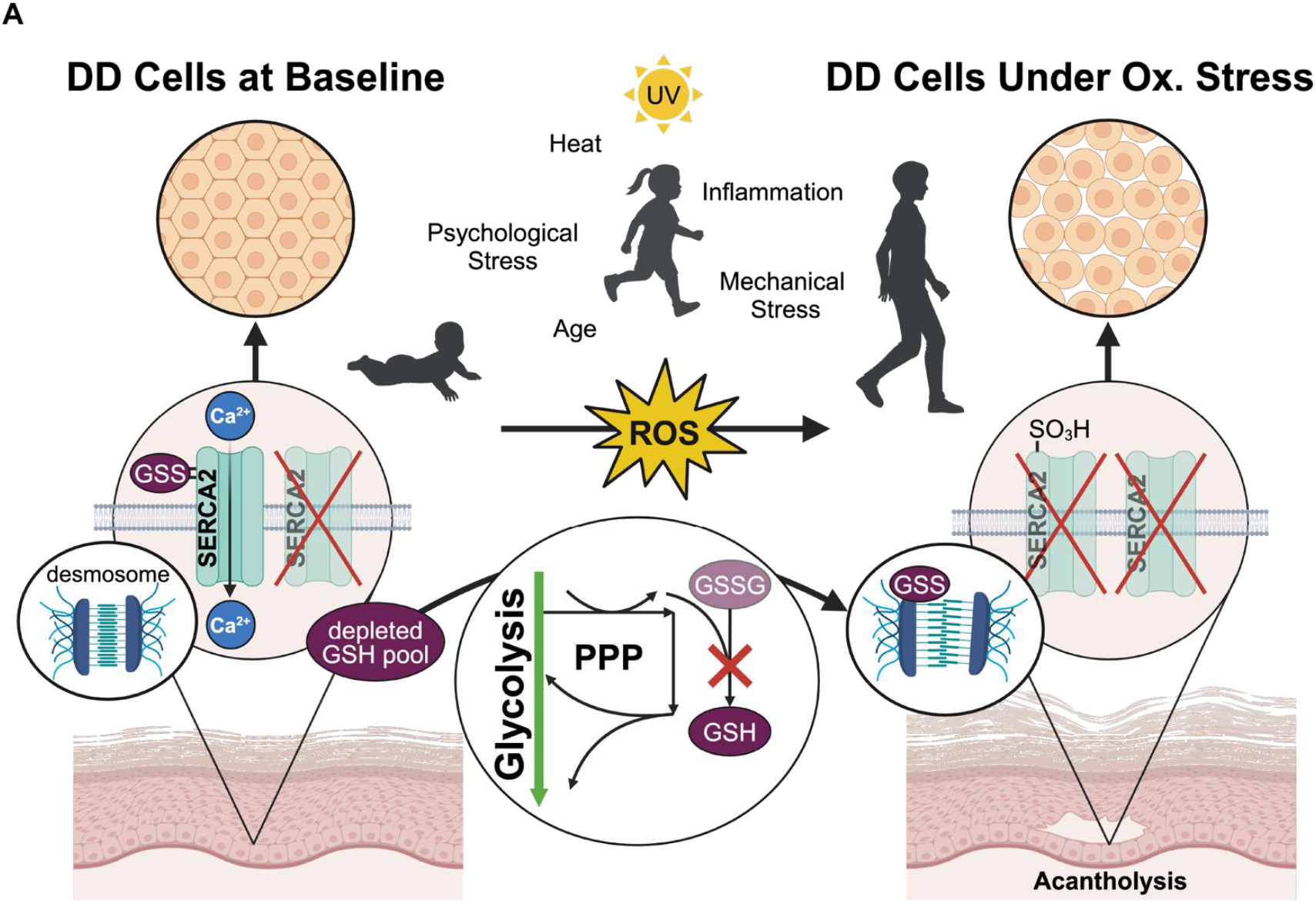
Model of stress induced acantholysis in DD. A schemiatic depicting the interplay of oxidative stress, cellular metabolism, and glutathione cycling in producing the phenotypic presentation of SERCA deficiency. DD cells at baseline have intact (though mildly weakened) desmosomal adhesions, increased glutathionylation of SERCA2 and a depleted pool of free glutathione. The addition of oxidative stress elicits a perturbed metabolic response, dissolution of adhesion and the perinuclear collapse of highly glutathionylated desmosomal components. Applying this model to patients, acantholytic regions would represent those that have inadequately responded to oxidative stress. Figure created using BioRender.com.

## Discussion

Genetic disorders linked to heterozygous variants can remain relatively hidden till later in life. Of those, a number are thought to represent haploinsufficiencies in which the mutant allele fails to produce expressed or active protein products. DD joins other haploinsufficiencies like the autoinflammatory condition, Bechet-like disease, possessing symptoms which ebb and flare in response to specific triggers. Based on metabolic and cell biological analysis of cells isolated from DD patients with heterozygous SERCA2 mutations, we propose a model of stress sensitivity hinging on the capacity of stress to ablate the remaining pool of active protein. This perspective may broadly apply to a number of haploinsufficiencies.

SERCA2-deficiency is commonly invoked as contributing to ER stress, a phenomenon characterized by misfolding of proteins in the ER. Indeed, transcriptomic analysis of SERCA2-depleted keratinocyte lines reveal a cell autonomous elevation of unfolded protein response pathways, in line with bulk RNA sequencing data of DD biopsies [19, 35]. Draining of SERCA-dependent ER calcium by agents like TG perturbs protein folding and, as such, represents a classic technique for initiating ER stress [36]. While treating the ER stress component of DD remains a rational approach, especially since trafficking of desmosomal cadherins from the ER is perturbed in DD [37], no such treatment has proven effective. ER stress, however, is a complex event that is integrated with a vast number of processes [38, 39]. For example, others have noted the capacity of menadione to elicit release of intracellular calcium stores [40, 41]. Understanding these intersections could illuminate orthogonal approaches. Those investigating depletion of the HHD-linked Golgi pump, ATP2C1, have successfully employed this strategy in linking Golgi stress, ROS and DNA damage repair defects [42, 43].

The misfolding which occurs when SERCA2-deficiency depletes luminal ER calcium likely contributes to heightened oxidative stress. Within the ER, improper formation of disulfide bonds elicits a repair response that entails breaking the incorrect bonds and initiating another round of disulfide bonding. The enzymatic reactions which catalyze this unfolding and refolding cycle both produce ROS and consume glutathione in the process [44]. Additional avenues link SERCA2-deficiency to ROS. Prior studies indicate that depletion of ER calcium stores increases mitochondrial calcium [45]. Enzymes involved in driving the citric acid cycle within mitochondria respond robustly to increased calcium levels, ultimately, overloading the electron transport chain and causing increased ROS [46]. Our understanding of ER-mitochondrial contacts, at sites referred to as MAMs (mitochondria associated membranes) continues to advance, shedding light on a potential route by which ER abnormalities lead to oxidative stress in DD cells [47]. This is a particularly intriguing perspective given recent work highlighting contact between the ER and desmosomes [48], setting up the possibility that all three organelles may influence each other through physical interactions. To what extent these processes contributes to oxidative stress sensitivity in our system awaits further investigation.

Among the most highly overexpressed transcripts in DD lesions are SLC7A11 and CHAC1 [35], both of which have been linked to ROS-induced cytotoxicity. CHAC1 enzymatically degrades glutathione, and its overexpression likely contributes to glutathione-deficiency in DD [49]. SLC7A11 exports glutamate, a precursor of glutathione, to import cystine, an oxidized precursor of cysteine. Elevated SLC7A11, in effect, floods the normally reducing cytoplasmic environment with the disulfide promoting activity of cystine whilst hampering production of protective glutathione [50]. This toxicity, referred to as disulfidoptosis entails improper disulfide bonding of the actin cytoskeleton. Given that serum-response factor (SRF) signaling, which directly responds to F-actin polymerization, is deficient in DD lesions [35], our current observations regarding sensitivity to oxidative stress could dovetail well with future examinations of cytoskeletal abnormalities in DD.

Beyond an earlier study highlighting sphingolipid abnormalities in SERCA2-deficient keratinocytes, few have addressed potential metabolite imbalances in DD [51]. Connections between the pentose phosphate pathway and glycolysis appear aberrant in DD cells, highlighting the need for future studies to address how calcium defects impinge on this relationship. Several potential avenues of investigation exist. Hexokinase converts glucose to glucose-6-phosphate (G6P), a metabolite which serves as the initial substrate for both glycolysis and the PPP, marking the first intersection of these parallel processes. Studies in the brain indicate that G6P binds and potently inhibits SERCA leading to an overload of cytoplasmic calcium and initiation of neuronal death [52]. The direct inhibition of SERCA by this metabolite could prove relevant to the sensitivity of DD cells to stressors, perhaps pushing the already compromised SERCA activity past a threshold from which cells cannot recover. Alternatively, the aberrantly high cytoplasmic calcium concentrations observed in DD cells could have significant impacts on enzymes within glycolysis and the PPP. A particularly interesting enzyme is transketolase (TKT), a calcium-responsive enzyme situated in the non-oxidative branch of the PPP responsible for conversions of ribose species [53]. One function of TKT is to convert excess ribose species in the PPP to glyceraldehyde-3-phosphate and feed it back into the lower steps of glycolysis to sustain pyruvate production. Regarding the present study, it is possible that elevated cytoplasmic calcium over-activates TKT and excessively shunts ribose back to glycolysis, accounting for blunted accumulation of R5P and failure to diminish pyruvate in DD cells exposed to menadione. Future studies should consider assessing the impact of TKT manipulation on DD phenotypes.

The rarity, phenotypic variability, and absence of an animal model have hampered efforts to develop novel treatments or repurpose existing interventions for DD. This study broadens the field of potential therapeutic targets by implicating the enzymatic machinery responsible for orchestrating a coordinated metabolic response to oxidative stress.

## Materials and Methods

### Donor Information, cell lines and culture conditions

We acquired DD keratinocyte cultures from patient biopsies collected at Northwestern University and Technion-Israel Institute of Technology. The study contains keratinocytes isolated from discarded abdominoplasty (CTL1) and lower back biopsies (CTL2-3) from three healthy adults. Cell isolates DD1 and DD2 were collected from the same patient carrying the pathogenic variant c.115A>G, p.Asn39Asp. CTL1-3 and DD1-2 were collected and isolated by the Northwestern Skin Biology & Diseases Resource-Based Center, under a protocol approved by the Northwestern University Institutional Review Board (IRB # STU00009443).

Cell lines DD3 and DD4 have been isolated from two distinct patients who both carry the pathogenic variant c.530A>C, p.Gln177Pro. Participants provided written informed consent prior to their inclusion in the study, according to the protocols approved by the Emek and Rabin Medical Centers (EMC-0086-15, RMC-0312-20) institutional review board.

Cell lines were maintained at 37°C in a humidified incubator with 5% CO2 on J2 feeder layers in FAD media [54]. DD3 and DD4 required the inclusion of 5uM Y27632 to establish and maintain cultures [55]. For consistency, all DD isolates and CTL1 and CTL2 were maintained in Y27632 unless specifically stated otherwise. For experiments, keratinocytes were removed from feeders and plated at 80-90% confluency in M154 (Thermo Fisher Scientific) growth media supplemented with 0.07 mM CaCl2, human keratinocyte growth supplement (HKGS; Thermo Fisher Scientific), gentamicin and amphotericin B 24-36 hours prior to the experiment. Y27632 was not present in experiments.

### Antibodies and Reagents

11-5F (gift from D. Garrod), anti-Glutathione conjugates (ViroGen, #101-A), K14, Y27632 (Cayman Chemical, #10005583), CDN1163 (Neurodon), NRDN501 (gift from Neurodon), n-acetylcysteine (Sigma, A7250), menadione (Sigma, M5625), anti-SERCA2 (ThermoFisher, MA3-919), mitomycin C (Sigma, M4287), Ponceau-S (Sigma,P7170)

### Western Blotting

Western blots were conducted by loading 20ug of lysate extracted with 1x Laemmli buffer+5% beta-mercaptoethanol. Samples were resolved on 7.5% SDS/PAGE gels purchased from BioRad and subjected to 80-100 volts in Tris/Glycine buffer containing 0.1%SDS. Protein was then transferred to nitrocellulose (0.45um pore size, Amersham) overnight at 22volts in Tris/Glycine buffer containing 20% methanol. After checking transfer quality with Ponceau S, blots were rinsed in TBS (pH 7.75, 200mM NaCl), blocked in 5% Milk/TBS for 1 hour, blotted overnight at 4C with anti-SERCA2 (2A7, 1:1000) in blocking buffer. Subsequently, blots were rinsed in TBS 3×10m and incubated with Goat-anti-mouse IgG-HRP conjugates (SantaCruz) for 30m. Following another 3×10m round of washing with TBS, blots were incubated with chemiluminescent solution for 2m (Millipore, Crescendo Substrate) and developed using a LiCor Odyssey instrument. For loading controls, blots were either blotted for GAPDH (Santa Cruz, 1:1000) or anti-vinculin (Sigma, 1:1000).

### Immunofluorescence and Microscopy

Cells plated in glass bottom 96-well plates were fixed using either ice-cold anhydrous methanol for 2 min or 4% paraformaldehyde (PFA) solution for 10 min at room temperature. Cells were blocked with 2% BSA and 1% normal goat serum, and those fixed with PFA were permeabilized using 0.5% Triton. All samples were mounted using a glycerol based mounting media as described here [56]. PLA analysis was performed as described in Hegazy et al [56] after fixation as described above. Fluorescent apotome images were acquired using a Zeiss Axio Observer, fitted with an X-Cite 120 LED Boost System, an Apotome.2 slide module, Axiocam 503 Mono digital camera, and a Plan-Apochromat 40x/1.4, Plan-Apochromat 63x/1.4 objective, Plan-Apochromat 100x/1.4 objective (Carl Zeiss). All images have undergone Orthogonal Projection in the Z plane using ZEN software.

### Dispase Assay

Cells were plated near confluency in 12-well dishes, then grown for 1-3 days in M154 until achieving a confluent monolayer. After washing with PBS, each well was treated with 2.4 U/mL of dispase (Sigma Millipore) diluted in high calcium M154 for 30 min at 37°C. Each lifted cell sheet was then placed in a 15 mL conical tube containing 3 mL of M154 and agitated in a rotator for 1 minute. Resulting fragments were returned to 12-well dishes, Zymo Research DNase was added to a final concentration of 1 unit in each well, and wells were imaged using a dissecting microscope (MZ6; Leica). Cell sheet fragments were counted manually in ImageJ from images.

### PLA and other FIJI analysis

Quantitation of western blot bands was conducted using ImageJ by measuring mean pixel intensity in regions of interest encompassing the band of a set size. After subtracting values obtained from blank portions of the blot and normalization to the loading control, relative protein abundance was calculated with respect to the control sample. PLA spots were counted in ImageJ using the protocol described here [57].

### Cellpose automated python analysis

We adapted a deep learning-based image segmentation framework Cellpose [24] and combined it with a series of quality control preprocessing steps to reliably obtain segmented metadata on hundreds of individual cells, similar to recent studies [25, 58]. Next, to quantify signal along the intercellular junctions, we formulated a new unbiased metric which we termed the Adhesion Stain Index, which goes along each cell’s outline and measures the number of pixels above a specified threshold and divide this number by the total number of pixels along the cell outline (**see Fig S2**).

### CellRox Assay

Examination of cellular ROS levels was conducted using CellRox dye (Invitrogen, C10444). Briefly, keratinocytes were plated in a tissue-culture treated 96-well plate with black walls in M154 with 0.07mM CaCl2 and incubated till confluency. Cells were then switched to 1.2mM CaCl2 overnight, pre-incubated with DMSO or 10uM CDN1163 for 1hr, loaded with CellRox dye in drug containing media for 30m, rinsed and replaced with fresh drug-containing media. An initial CellRox fluorescence reading was taken, then menadione was added to 20uM and reading were taken every 60m.

### Metabolite sample preparation and analysis

Cells were grown in 10cm dishes in M154 (0.07mM CaCl2) till 75% confluent. At that point, cells were switched to 1.2mM CaCl2 with 10uM menadione or vehicle (EtOH) for 4hrs. Cells were rinsed twice with ice cold PBS, then lysed on dry ice with 80% methanol pre-chilled to –80C. After scraping into microfuge tubes, lysates were subjected to three rounds of freezing and thawing in liquid nitrogen, followed by incubation overnight at –20C. Lysates were spun at maximum speed at 4C for 15m to pellet precipitated protein. Supernatant was removed, dried via SpeedVac, and delivered to the Northwestern University Metabolomics Core for LC/MS analysis with injections normalized to the concentration of resolubilized protein pellets. Raw peak intensities were then analyzed via Metaboanalyst 6.0.

For unpaired analysis (**Figure 3A-B**), metabolites were collected for three independent experiments comparing one control (CTL1) and the four DD (DD1-4) isolates. In addition, three experiments were conducted with two control patient isolates (CTL1-2) and two DD isolates (DD1-2) maintained without the Y27631 used to prolong culture lifespan. In total, 27 samples were subjected to comparison. Of 298 potentially identifiable metabolites, 145 passed the quality control filters available through Metaboanalyst 6.0 including removal of the least abundant 20% and 10% demonstrating near-constant values via interquartile range. Samples were normalized to the median peak value of each metabolite prior to statistical analysis.

For pairwise comparisons, the response of each metabolite within each isolate to menadione was measured per experimental repeat following normalization to median peak intensity within the three biological repeats for CTL1 and DD1-4. Correlation maps represent the average response of CTL1 and the DD isolates to menadione (**Figure 3C**). For comparison of individual metabolites across all conditions (**Figure 3D-F**), each metabolite for CTL1 and DD1-4 was normalized to sample peak intensity medians in Metaboanalyst 6.0.

### GSH Plate Reader Assay

Keratinocytes were plated in a 6-well dish (1 million cells per well) in M154 (0.07mM CaCl2) and incubated till 80-100% confluency. Cells were switched to 1.2mM CaCl2 +/- 10uM menadione for 4hrs. Cells were trypsinized, quenched, rinsed with PBS, pelleted and spun down. Cells were resuspended in 200ul PBS then lysed by freezing and thawing in liquid nitrogen three times, spinning at max speed in microfuge for 15m at 4C. 150ul of supernatant was collected and frozen. Pellet was also frozen for quantification of cell debris pellet for normalization. Supernatant was subjected to a fluorometric glutathione assay (AnaSpec, AS-72158).

### Statistical analysis and data plotting

Plots were generated using Graphpad Prism 10 and Adobe Illustrator. All statistics were performed on raw data to preserve variability of CTL samples, although some plots display fold change for visual clarity. Significances were determined via One-way Anova or Two-tailed T-tests, except where indicated. Non significance is indicated by ‘ns’; p-values less than 0.05 are indicated with one star, p-values less than 0.01 are indicated with 2 stars, and p-values less than 0.001are indicated with three stars.

## Acknowledgements

The authors gratefully acknowledge a generous donation from the Lee family, who directly supported this work via funding to KJG, EFM, LG and RMH. KJG acknowledges support from National Institutes of Health grants R01AR041836, R01AR043380, and RO1CA228196; a LEO Foundation grant; and the JL Mayberry endowment. MP and AS acknowledge support from NIH grants R01NS057499 and R35NS132349. ASP acknowledges support from the National Institute of Arthritis and Musculoskeletal and Skin Diseases R01AR44619 and R01AR068375 (ASP). The authors would also like to acknowledge support from the Northwestern University Skin Biology and Disease Resource-Based Center P30AR075049. CLS acknowledges grant support from National Institutes of Health R03AR082896, R03TR005428, the LEO Foundation (LF-OC-23-001393), the Foundation for Ichthyosis and Related Skin Types. EB acknowledges support by the Leo foundation grant (LF-OC-20-000585). Metabolomics experiments were performed by Metabolomics Core Facility at Robert H. Lurie Comprehensive Cancer Center (generously supported by NCI CCSG P30 CA060553) of Northwestern University.

