## Supplemental Figures S1-3 for "Failed Metabolic Adaptation to Stress Underlies Pathogenesis in a Heterozygous Genetic Disorder"

**Supplemental Figure Legends:**

**Figure S1. Calcium imaging data.** A) Ratiometric Fura-2 signal traces indicating cytoplasmic calcium concentrations from three independent experiments, n1-3. Mean trace values +/- s.e.m. at each time point of the experiment derived from 50-70 individual cells from imaging of three separate plates of CTL1 and DD2 isolates. B) Baseline differences in cytoplasmic calcium levels derived from data in panel A. Small dots represent values from individual cells, large open shapes represent the mean value from each experiment. Asterix indicate $p<0.05$ of two-tailed paired t-test of the mean values. C) SOCE quantitation. Small dots represent values from individual cells, large open shapes represent the mean value from each experiment. p-values derived from paired t-tests. A.U.C= area under the curve.

**Figure S2. Cellpose-based quantitation of cell border localization.** a) Examples of Cellpose-based segmentation of cell nuclei and borders. A cell is defined by an Adhesion Stain mask, inside the bounds of which a Nuclei Stain mask is contained. Left panels represent computational output of cell bodies and nuclei, center panels represent merge of computed masks with corresponding immunofluorescent and nuclei stains and the right panels display outlined cell boundaries (red) and nuclei boundaries upon which subsequent measurements are based. b) Workflow for determining adhesion stain index of a given marker (e.g. DP). c) Representative quantitation from one experiment demonstrating the adhesion staining index values obtained from >200 individual cells for CTL and DD cells treated with either DMSO or CDN1163 (10uM). Dots represent values from each cell, box =25-75$^{th}$ percentile, line=median. The mean of such a distribution--for each condition of each experiment--is what is displayed in Figures 1, 2 and 5. Examples of individual cells across the spectrum of adhesion stain indexes stained for DP are displayed. d) The impact of varying the pixel intensity threshold on adhesion stain index values. Dotted line represents threshold chosen for this study.

**Figure S3. Additional metabolomics information.** a) Partial least squares discriminative analysis (PLS-DA) of DD cell isolates versus a control keratinocyte isolate (CTL2). Each dot represents the metabolite profile of a given isolate for one of three independent experiments. b) Volcano plots of individual DD isolate metabolites compared to a control keratinocyte isolate (CTL2). Data represent values calculated from 3 independent experiments after normalization to median peak values for each sample. Log2 fold change (FC) derived from mean peak values, p-values determined from unpaired t-tests of those means. Values for R5P and S7P are denoted. c) List of glycolysis-TCA intermediates coordinately downregulated upon menadione treatment of CTL2 as well as a list of anti-correlated PPP intermediates. d) Measurement of reduced glutathione (GSH) levels via plate reader assay. Asterix indicate statistical significance as calculated via a one-sample t-test (hypothetical mean =1).

**Figure S1:**

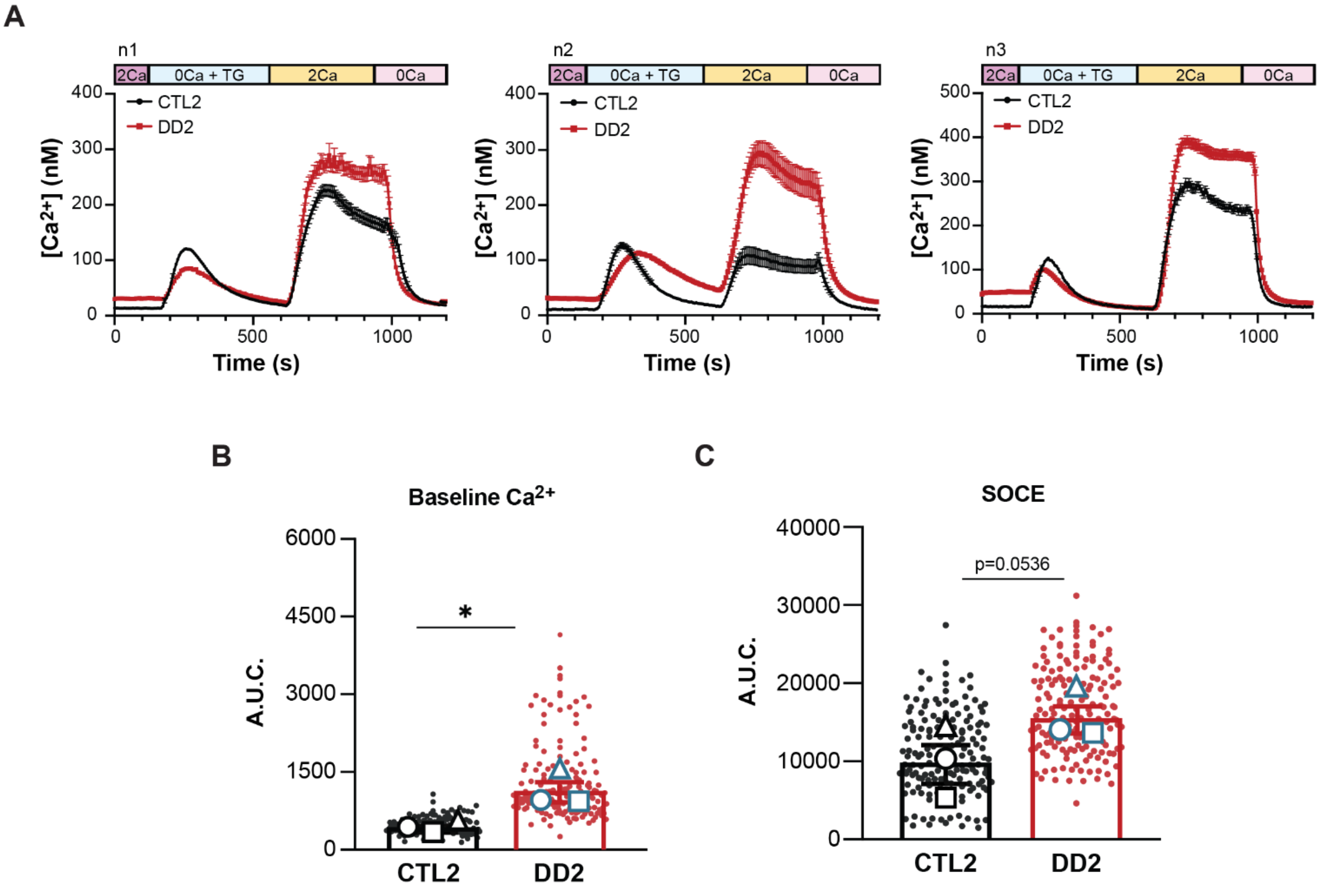
A
n1
2Ca
0Ca + TG
2Ca
0Ca
CTL2
DD2
[Ca2+] (nM)
Time (s)
n2
n3
B
Baseline Ca2+
A.U.C.
*
CTL2
DD2
C
SOCE
p=0.0536

**Figure S2:**

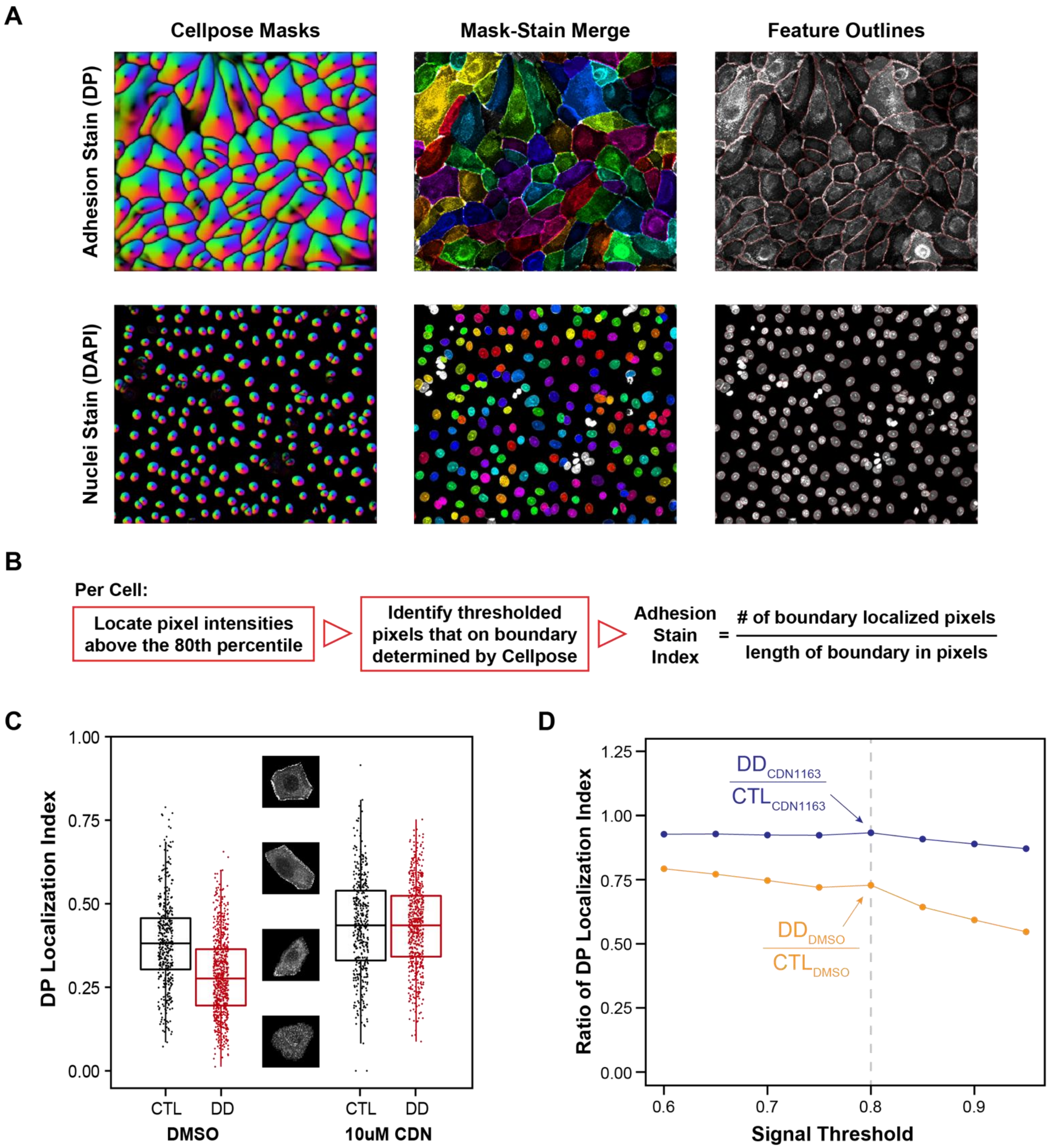
A
Cellpose Masks
Mask-Stain Merge
Feature Outlines
Adhesion Stain (DP)
Nuclei Stain (DAPI)
B
Per Cell:
Locate pixel intensities above the 80th percentile
Identify thresholded pixels that on boundary determined by Cellpose
Adhesion Stain Index = # of boundary localized pixels / length of boundary in pixels
C
DP Localization Index
1.00
0.75
0.50
0.25
0.00
CTL
DD
CTL
DD
DMSO
10uM CDN
D
Ratio of DP Localization Index
1.25
1.00
0.75
0.50
0.25
0.00
DD CDN1163 / CTL CDN1163
DD DMSO / CTL DMSO
0.6
0.7
0.8
0.9
Signal Threshold

**Figure S3:**

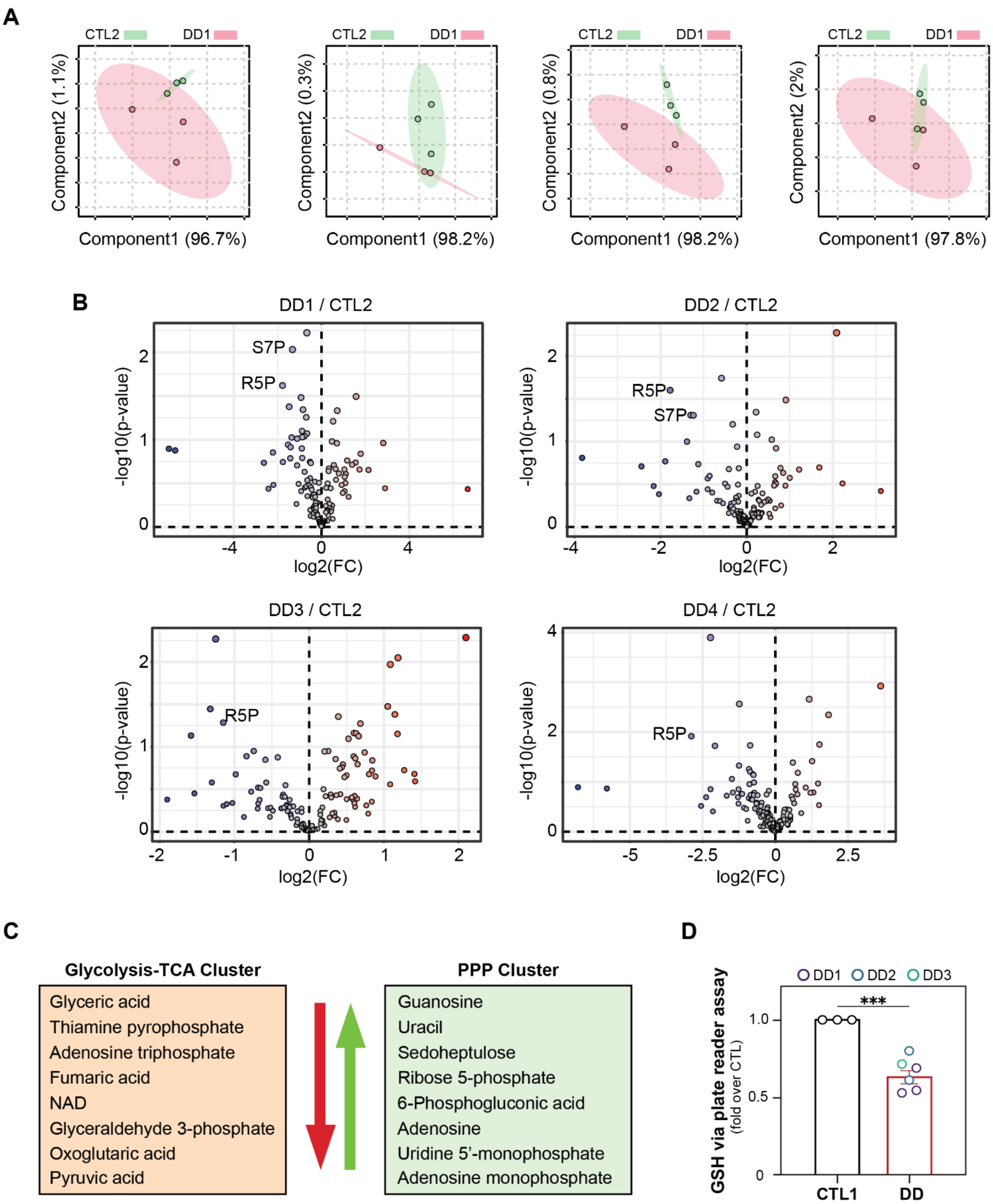
A
CTL2
DD1
Component2 (1.1%)
Component1 (96.7%)
Component2 (0.3%)
Component1 (98.2%)
Component2 (0.8%)
Component1 (98.2%)
Component2 (2%)
Component1 (97.8%)
B
DD1 / CTL2
S7P
R5P
DD2 / CTL2
R5P
S7P
DD3 / CTL2
R5P
DD4 / CTL2
R5P
-log10(p-value)
log2(FC)
C
Glycolysis-TCA Cluster
Glyceric acid
Thiamine pyrophosphate
Adenosine triphosphate
Fumaric acid
NAD
Glyceraldehyde 3-phosphate
Oxoglutaric acid
Pyruvic acid
PPP Cluster
Guanosine
Uracil
Sedoheptulose
Ribose 5-phosphate
6-Phosphogluconic acid
Adenosine
Uridine 5'-monophosphate
Adenosine monophosphate
D
DD1 DD2 DD3
GSH via plate reader assay
(fold over CTL)
***
CTL1
DD
